## supplementary tables and figures for "Reactive oxygen species trigger downward vertical migration in diatom microphytobenthic biofilms as a strategy to cope with oxidative stress"

### SUPPLEMENTARY FIGURES AND TABLES

***Table s1.*** *Equations adapted from Consalvey (2005), and Christof Klughammer and Ulrich Schreiber (2008) used to calculate photosynthetic parameters.*

| Photosynthetic parameter | Equation | Description |
|---|---|---|
| **α-slope** (arbitrary unit) | $\frac{1}{c}$ | Maximum light use efficiency of photosystem II at low irradiance, derived from the initial slope of the photosynthesis-irradiance (P-E) curve.<br>*c : empirical coefficient derived from model fitting.* |
| **Fv/Fm** (unitless) | $\frac{Fm - F0}{Fm}$ | Maximum quantum yield of photosystem II photochemistry under dark-adapted state.<br>*Fm Maximum fluorescence yield of dark-adapted sample; $F_0$ Minimum fluorescence yield of dark-adapted sample.*<br>*Under actinic illumination, this calculation is replaced by Fm'-F/Fm', which corresponds to the effective photosystem II efficiency during subsequent steps of the light curve.* |
| **rETR** (arbitrary unit) | $\frac{Fm' - F}{Fm'} \times \frac{Irradiance}{2}$ | Relative electron transport rate of photosystem II, with the term $rETR_{m\ obs}$ referring to the maximum rETR observed in the light curve measured without extrapolation.<br>*Fm' Maximum fluorescence of illuminated sample; F Fluorescence yield in the light-adapted state measured briefly before application of a Saturation Pulse; Irradiance refers to the incident light μmol photons $m^{-2}$ $s^{-1}$.* |
| **NPQ** (unitless) | $\frac{Fm - Fm'}{Fm'}$ | Non-photochemical quenching, with the term $NPQ^{1000}$ referring to the maximum NPQ value obtained after the final one-minute light curve step at 1000 μmol photons $m^{-2}$ $s^{-1}$, followed by the saturating flash used to determine Fm′. |
| **Y(NPQ)** (unitless) | $\frac{F}{Fm'} - \frac{F}{Fm}$ | Regulated non-photochemical quenching, with the term $Y(NPQ)^{1000}$ referring to the maximum Y(NPQ) value obtained after the final one-minute light curve step at 1000 μmol photons $m^{-2}$ $s^{-1}$, followed by the saturating flash used to determine Fm′. |
| **Y(NO)** (unitless) | $\frac{F}{Fm}$ | Non-regulated non-photochemical quenching, with the term $Y(NO)_{m\ obs}$ referring to the maximum Y(NO) observed within the measured light curve range (no extrapolation). |

***Table s2**. Pigment diversity in microphytobenthic samples, with an example of a chromatogram.*

| Pigment | λ Integration (nm) | Retention time (min) |
|---|---|---|
| Chlorophyll c2 (Chlc2) | 446 | 9.5 |
| Fucoxanthin like 1 (F-l1) | 448 | 11.2 |
| Fucoxanthin (F) | 448 | 12.2 |
| Fucoxanthin like 2 (F-l2) | 448 | 12.6 |
| Fucoxanthin like 3 (F-l3) | 448 | 12.9 |
| Fucoxanthin like 4 (F-l4) | 448 | 13.1 |
| Fucoxanthin like 5 (F-l5) | 448 | 13.53 |
| Neoxanthin (N) | 441 | 13.85 |
| Pheophorbide a 1 (Pda1) | 663 | 14.12 |
| Violaxanthin (V) | 441 | 14.25 |
| Pheophorbide a 2 (Pda2) | 663 | 14.40 |
| Fucoxanthin like 6 (F-l6) | 448 | 14.88 |
| Diadinoxanthin (Dd) | 448 | 15.31 |
| Fucoxanthin like 7 (F-l7) | 448 | 15.77 |
| Antheraxanthin (A) | 448 | 16.13 |
| Unknown Carotenoid 1 (Uc1) | 448 | 16.38 |
| Unknown Carotenoid 2 (Uc2) | 448 | 16.58 |
| Unknown Carotenoid 3 (Uc3) | 448 | 16.9 |
| Diatoxanthin (Dt) | 454 | 17.17 |
| Unknown Carotenoid 4 (Uc4) | 448 | 17.43 |
| Pheophorbide a 3 (Pda3) | 663 | 17.63 |
| Lutein (L) | 448 | 17.68 |
| Zéaxanthin (Z) | 448 | 17.96 |
| Unknown Carotenoid 5 (Uc5) | 448 | 18.2 |
| Unknown Carotenoid 6 (Uc6) | 448 | 18.47 |
| Unknown Carotenoid 7 (Uc7) | 448 | 18.70 |
| Unknown Carotenoid 8 (Uc8) | 448 | 19.03 |
| Unknown Carotenoid 9 (Uc9) | 448 | 21.35 |
| Chlorophyll a like 1 (Chla-l1) | 412 | 21.56 |
| Chlorophyll a like 2 (Chla-l2) | 431 | 22.2 |
| Chlorophyll a like 3 (Chla-l3) | 412 | 22.45 |
| Chlorophyll a allomer (Chla-allomer) | 431 | 23 |
| Chlorophyll a (Chla) | 431 | 23.6 |
| Chlorophyll a epimer (Chla-epimer) | 431 | 24.1 |
| Chlorophyll a like 4 (Chla-l4) | 431 | 24.32 |
| Chlorophyll a like 5 (Chla-l5) | 431 | 24.65 |
| Chlorophyll a like 6 (Chla-l6) | 431 | 25.3 |
| Pheophytin a like 1 (Phal1) | 663 | 26.1 |
| Chlorophyll a like 7 (Chla-l7) | 431 | 26.45 |
| Pheophytin a (Pha) | 663 | 26.6 |
| Pheophytin a like 2 (Phal2) | 663 | 26.8 |
| β-ε-carotene (βε) | 454 | 27.4 |
| β-β-carotene (ββ) | 454 | 27.7 |
| Pyropheophytin a (Pya) | 663 | 27.8 |

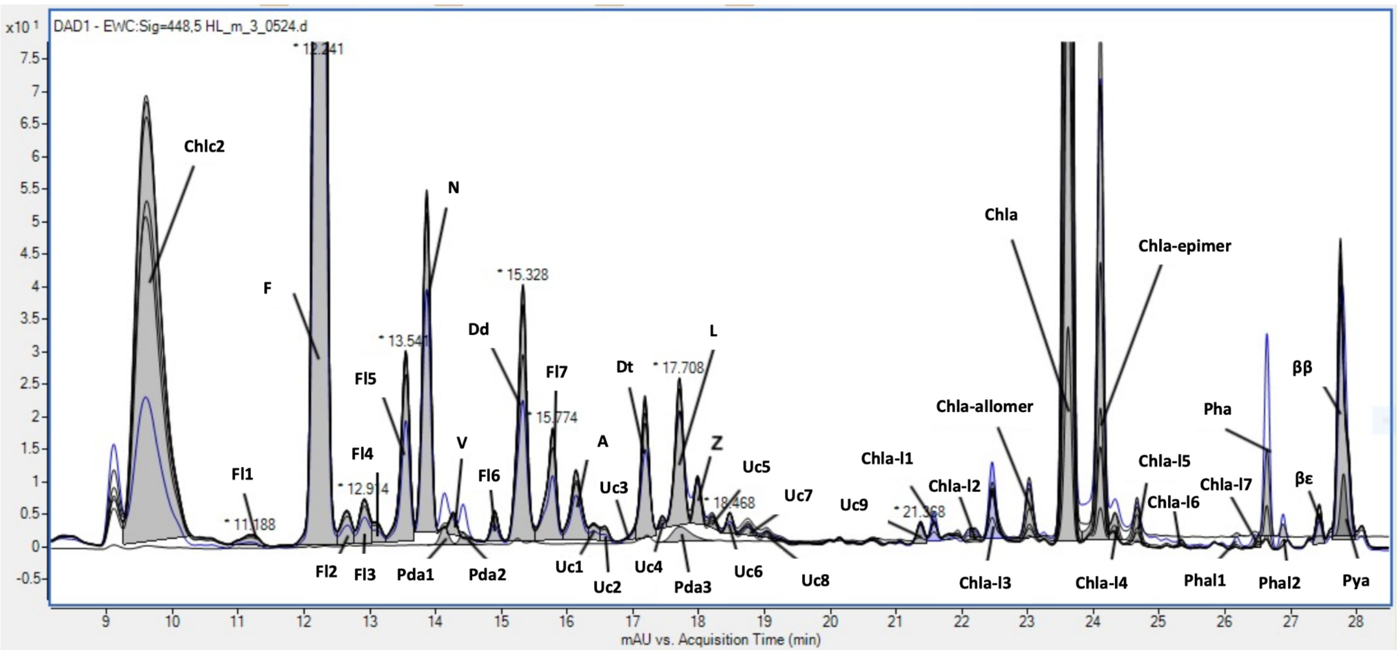

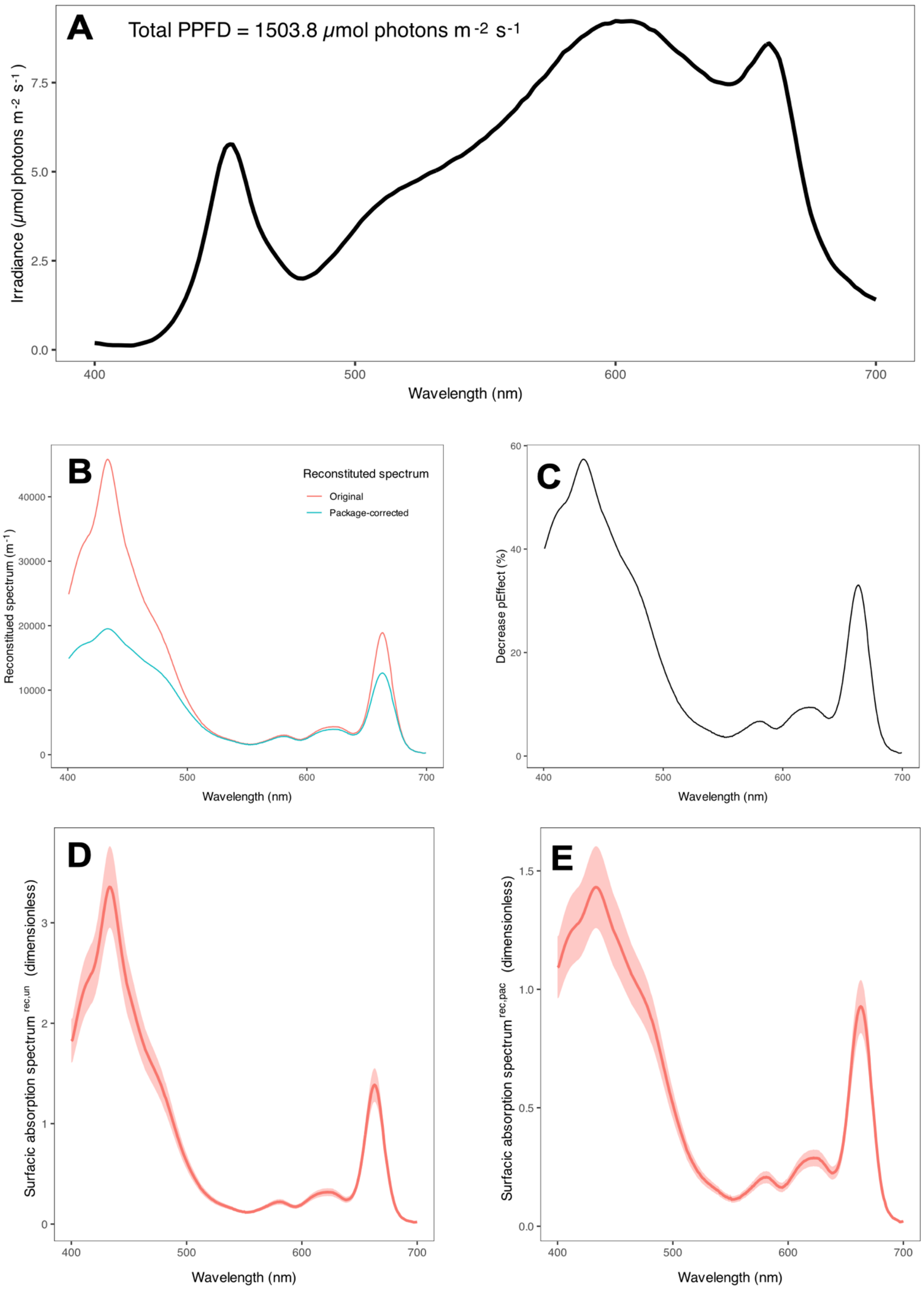


***Figure s1.*** *Qphar calculation and package effect correction.* ***A.*** *Spectrum of incident irradiance;* ***B.*** *Original and package-corrected, average reconstructed absorption spectrum of sediment-free biofilm community (dark-adapted samples);* ***C.*** *Average absorbance loss (%) for dark-adapted sediment-free biofilm samples due to the package effect;* ***D and E.*** *Variability in the reconstructed absorption spectrum of replicates (n = 5) of dark-adapted sediment-free biofilm samples, without package effect correction and with package effect correction, respectively.*

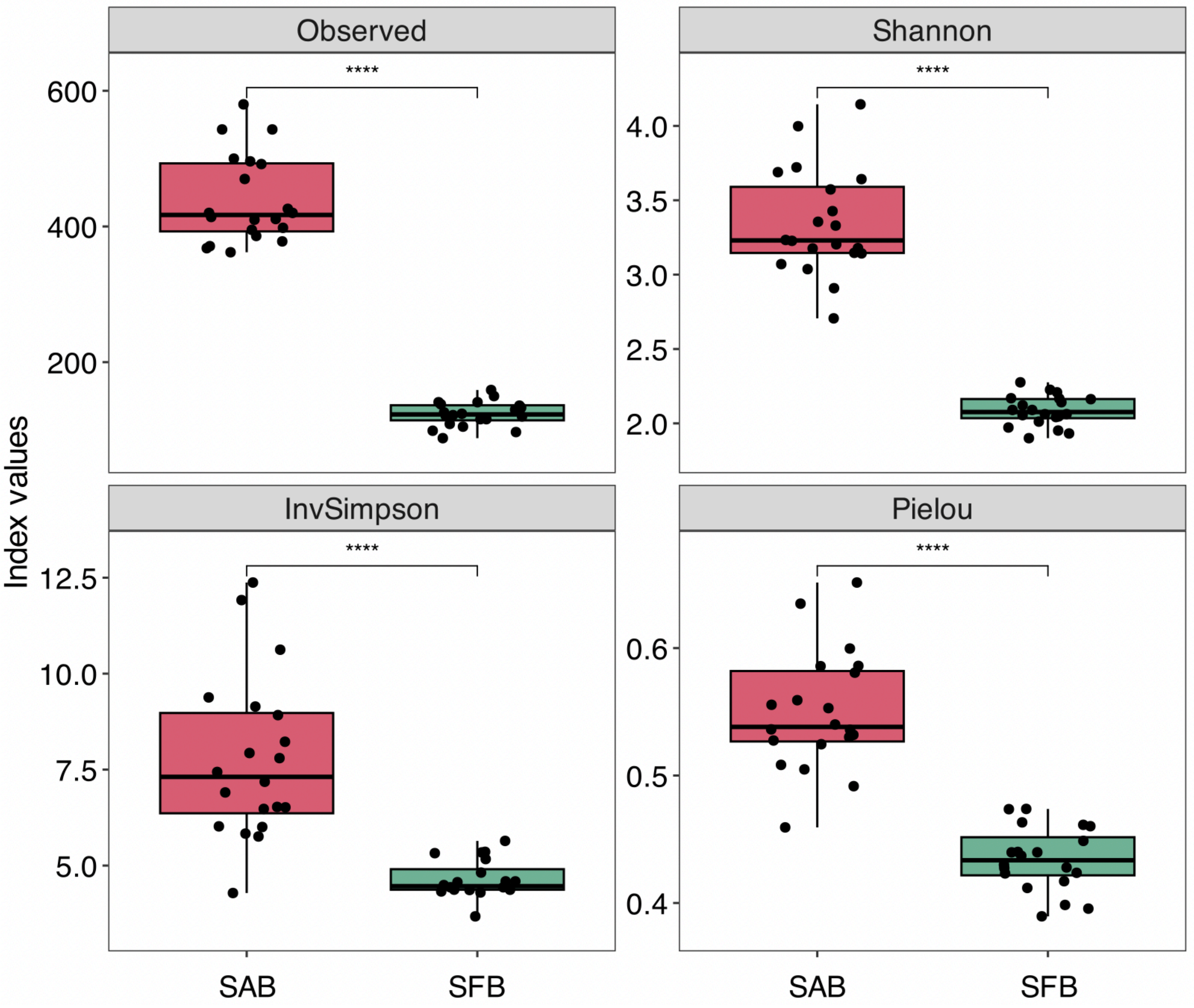


***Figure s2.*** *Boxplot of different diversity indices for eukaryotic community based on Sediment-Associated Biofilm (SAB) and Sediment-Free Biofilm samples (SFB) (statistically tested by Wilcoxon-Mann-Whitney; n = 20, P < 0.0001****).*

***Table s3.*** *Table of raw photosynthetic parameters values (mean ± SD) before and after the different treatments: High-Light (HL), Hydrogen Peroxide (HP), and Cold Atmospheric Plasma (PL), for both Sediment-Associated Biofilm (SAB) and Sediment-Free Biofilm (SFB) communities. Dark-Adapted control (DA)(1), (2), and (3) correspond to samples measured in parallel to the HL, HP, and PL experimental treatments, respectively. α-slope (arbitrary unit) corresponds to the maximum light use efficiency of photosystem II under low irradiances. Fv/Fm (unitless) corresponds to the maximum quantum yield of photosystem II photochemistry under dark-adapted state. $rETR_{m\ obs}$ (arbitrary unit) and $Y(NO)_{m\ obs}$ (unitless) refer to the maximum values observed within the measured light curve range (without extrapolation). $NPQ^{1000}$ (unitless) and $Y(NPQ)^{1000}$ (unitless) correspond to the maximum values measured after the final one-minute light curve step at 1000 µmol photons $m^{-2}\ s^{-1}$ (without extrapolation). $F_0$ refers to the minimum dark-adapted fluorescence yield (arbitrary unit). Eopt refers to the optimal light parameter for autotrophic cells (µmol photons $m^{-2}\ s^{-1}$). Ek refers to the light saturation coefficient where the cell begins to activate photoprotective mechanisms (µmol photons $m^{-2}\ s^{-1}$). All fluorescence measurements were performed on dark-adapted samples. For each photophysiological parameter, the mean was calculated from triplicate values (n = 3).*

| Treatment | α-slope | Fv/Fm | rETRm obs | NPQ1000 | YNPQ1000 | YNOm obs | F0 | Eopt | Ek |
|---|---|---|---|---|---|---|---|---|---|
| **DA SAB (1) Before** | 0.35 (0.01) | 0.70 (0.00) | 85.04 (3.70) | 1.72 (0.16) | 0.52 (0.02) | 0.42 (0.02) | 3045.33 (473.50) | 876.79 (97.84) | 246.32 (4.27) |
| **DA SAB (1) After** | 0.34 (0.00) | 0.69 (0.01) | 87.47 (2.97) | 1.42 (0.27) | 0.48 (0.04) | 0.45 (0.03) | 2323.00 (330.59) | 875.97 (42.19) | 261.31 (7.23) |
| **HL SAB Before** | 0.35 (0.01) | 0.69 (0.01) | 84.19 (1.82) | 1.56 (0.17) | 0.51 (0.02) | 0.44 (0.02) | 3045.33 (431.86) | 862.04 (30.53) | 246.15 (7.78) |
| **HL SAB After** | 0.25 (0.03) | 0.53 (0.03) | 67.87 (7.99) | 2.13 (0.20) | 0.59 (0.01) | 0.49 (0.02) | 1542.00 (180.91) | 743.48 (53.19) | 283.67 (7.70) |
| **DA SFB (1) Before** | 0.32 (0.02) | 0.67 (0.01) | 70.56 (1.53) | 2.69 (0.17) | 0.63 (0.01) | 0.35 (0.01) | 1253.00 (135.97) | 697.26 (116.93) | 219.95 (16.18) |
| **DA SFB (1) After** | 0.34 (0.01) | 0.69 (0.01) | 60.82 (5.93) | 2.11 (0.09) | 0.60 (0.00) | 0.39 (0.01) | 1229.33 (118.46) | 629.67 (31.42) | 179.23 (21.43) |
| **HL SFB Before** | 0.32 (0.01) | 0.67 (0.01) | 74.43 (3.94) | 2.73 (0.18) | 0.62 (0.02) | 0.35 (0.00) | 1139.67 (132.59) | 822.14 (41.08) | 234.91 (15.73) |
| **HL SFB After** | 0.26 (0.03) | 0.53 (0.04) | 47.07 (9.11) | 2.84 (0.31) | 0.67 (0.01) | 0.49 (0.06) | 1632.00 (132.84) | 602.96 (70.22) | 182.49 (19.04) |
| **DA SAB (2) Before** | 0.35 (0.01) | 0.70 (0.01) | 90.85 (2.60) | 0.74 (0.49) | 0.32 (0.17) | 0.62 (0.19) | 2128.33 (649.98) | 925.47 (61.98) | 259.57 (9.69) |
| **DA SAB (2) After** | 0.34 (0.01) | 0.71 (0.01) | 87.59 (4.59) | 0.96 (0.10) | 0.40 (0.02) | 0.52 (0.03) | 2740.00 (425.97) | 805.90 (107.60) | 260.05 (9.29) |
| **HP SAB Before** | 0.35 (0.00) | 0.71 (0.00) | 89.51 (5.97) | 1.12 (0.04) | 0.43 (0.01) | 0.50 (0.00) | 2127.00 (114.53) | 932.51 (65.64) | 256.23 (16.94) |
| **HP SAB After** | 0.32 (0.01) | 0.65 (0.01) | 93.56 (2.16) | 0.02 (0.04) | 0.02 (0.03) | 0.86 (0.06) | 573.67 (59.94) | 997.74 (21.54) | 289.41 (5.95) |
| **DA SFB (2) Before** | 0.34 (0.00) | 0.69 (0.01) | 78.06 (1.39) | 2.66 (0.09) | 0.61 (0.00) | 0.32 (0.01) | 1445.67 (105.45) | 774.52 (108.62) | 234.21 (5.04) |
| **DA SFB (2) After** | 0.34 (0.01) | 0.69 (0.01) | 70.57 (8.39) | 2.27 (0.26) | 0.60 (0.03) | 0.36 (0.01) | 1266.00 (79.37) | 718.08 (90.44) | 211.62 (33.60) |
| **HP SFB Before** | 0.32 (0.01) | 0.69 (0.00) | 73.75 (4.80) | 2.38 (0.07) | 0.61 (0.01) | 0.34 (0.01) | 1276.00 (105.36) | 653.55 (60.50) | 229.40 (19.40) |
| **HP SFB After** | 0.34 (0.02) | 0.68 (0.01) | 68.86 (7.36) | 2.46 (0.22) | 0.62 (0.02) | 0.35 (0.01) | 1358.67 (109.13) | 699.40 (160.25) | 205.03 (27.30) |
| **DA SAB (3) Before** | 0.34 (0.01) | 0.68 (0.01) | 88.67 (3.38) | 1.46 (0.16) | 0.49 (0.02) | 0.49 (0.02) | 1638.33 (49.33) | 911.13 (40.12) | 266.01 (15.28) |
| **DA SAB (3) After** | 0.34 (0.01) | 0.67 (0.01) | 89.12 (5.37) | 1.10 (0.45) | 0.42 (0.08) | 0.56 (0.09) | 1701.67 (314.33) | 915.32 (34.48) | 267.38 (16.63) |
| **PL SAB Before** | 0.34 (0.01) | 0.67 (0.02) | 85.16 (1.78) | 1.20 (0.31) | 0.45 (0.06) | 0.55 (0.06) | 1296.00 (234.16) | 943.36 (65.61) | 254.74 (9.17) |
| **PL SAB After** | 0.23 (0.01) | 0.57 (0.01) | 66.10 (1.71) | 0.39 (0.04) | 0.24 (0.02) | 0.63 (0.02) | 357.33 (32.96) | 906.42 (51.02) | 286.90 (7.73) |
| **DA SFB (3) Before** | 0.33 (0.01) | 0.70 (0.01) | 71.89 (3.09) | 2.97 (0.17) | 0.65 (0.00) | 0.34 (0.02) | 1625.67 (148.76) | 639.81 (27.66) | 218.65 (9.28) |
| **DA SFB (3) After** | 0.34 (0.02) | 0.69 (0.01) | 70.19 (6.33) | 2.51 (0.18) | 0.62 (0.00) | 0.37 (0.02) | 1432.00 (218.47) | 734.44 (151.18) | 209.42 (21.94) |
| **PL SFB Before** | 0.35 (0.02) | 0.69 (0.02) | 68.75 (9.49) | 2.34 (0.44) | 0.60 (0.02) | 0.40 (0.06) | 1552.00 (351.51) | 813.73 (370.73) | 196.06 (18.42) |
| **PL SFB After** | 0.33 (0.01) | 0.66 (0.00) | 53.55 (8.27) | 2.15 (0.01) | 0.62 (0.01) | 0.37 (0.01) | 1312.67 (35.12) | 572.27 (120.11) | 164.80 (21.01) |

***Table s4.*** *Table of average changes (%) in photosynthetic parameters induced by treatments: High-Light (HL), Hydrogen Peroxide (HP), and Cold Atmospheric Plasma (PL), for both Sediment-Associated Biofilm (SAB) and Sediment-Free Biofilm (SFB) communities. Dark-Adapted control (DA) (1), (2), and (3) correspond to samples measured in parallel to the HL, HP, and PL experimental treatments, respectively. The average values are calculated from the raw photosynthetic parameter data (supplementary Table 3) using the following equation:* $((\bar{x}_{After\ treatment} - \bar{x}_{Before\ treatment})/\bar{x}_{Before\ treatment}) x100$. *α-slope (arbitrary unit) corresponds to the maximum light use efficiency of photosystem II under low irradiances. Fv/Fm (unitless) corresponds to the maximum quantum yield of photosystem II photochemistry under dark-adapted state.* $rETR_{m\ obs}$ *(arbitrary unit) and* $Y(NO)_{m\ obs}$ *(unitless) refer to the maximum values observed within the measured light curve range (without extrapolation).* $NPQ^{1000}$ *(unitless) and* $Y(NPQ)^{1000}$ *(unitless) correspond to the maximum values measured after the final one-minute light curve step at 1000 µmol photons* $m^{-2}$ $s^{-1}$ *(without extrapolation).* $F_0$ *refers to the minimum dark-adapted fluorescence yield (arbitrary unit). Eopt refers to the optimal light parameter for autotrophic cells (µmol photons* $m^{-2}$ $s^{-1}$*). Ek refers to the light saturation coefficient where the cell begins to activate photoprotective mechanisms (µmol photons* $m^{-2}$ $s^{-1}$*). All fluorescence measurements were performed on dark-adapted samples.*

| Treatment | α-slope | Fv/Fm | rETRm obs | NPQ1000 | YNPQ1000 | YNOm obs | F0 | Eopt | Ek |
|---|---|---|---|---|---|---|---|---|---|
| **DA SAB (1)** | -2.86 | -1.43 | 2.86 | -17.44 | -7.69 | 7.14 | -23.72 | -0.09 | 6.09 |
| **HL SAB** | -28.57 | -23.19 | -19.38 | 36.54 | 15.69 | 11.36 | -49.37 | -13.75 | 15.24 |
| **DA SFB (1)** | 6.25 | 2.99 | -13.80 | -21.56 | -4.76 | 11.43 | -1.89 | -9.69 | -18.51 |
| **HL SFB** | -18.75 | -20.90 | -36.76 | 4.03 | 8.06 | 40.00 | 43.20 | -26.66 | -22.31 |
| **DA SAB (2)** | -2.86 | 1.43 | -3.59 | 29.73 | 25.00 | -16.13 | 28.74 | -12.92 | 0.18 |
| **HP SAB** | -8.57 | -8.45 | 4.52 | -98.21 | -95.35 | 72.00 | -73.03 | 7.00 | 12.95 |
| **DA SFB (2)** | 0.00 | 0.00 | -9.60 | -14.66 | -1.64 | 12.50 | -12.43 | -7.29 | -9.65 |
| **HP SFB** | 6.25 | -1.45 | -6.63 | 3.36 | 1.64 | 2.94 | 6.48 | 7.02 | -10.62 |
| **DA SAB (3)** | 0.00 | -1.47 | 0.51 | -24.66 | -14.29 | 14.29 | 3.87 | 0.46 | 0.52 |
| **PL SAB** | -32.35 | -14.93 | -22.38 | -67.50 | -46.67 | 14.55 | -72.43 | -3.92 | 12.62 |
| **DA SFB (3)** | 3.03 | -1.43 | -2.36 | -15.49 | -4.62 | 8.82 | -11.91 | 14.79 | -4.22 |
| **PL SFB** | -5.71 | -4.35 | -22.11 | -8.12 | 3.33 | -7.50 | -15.42 | -29.67 | -15.94 |

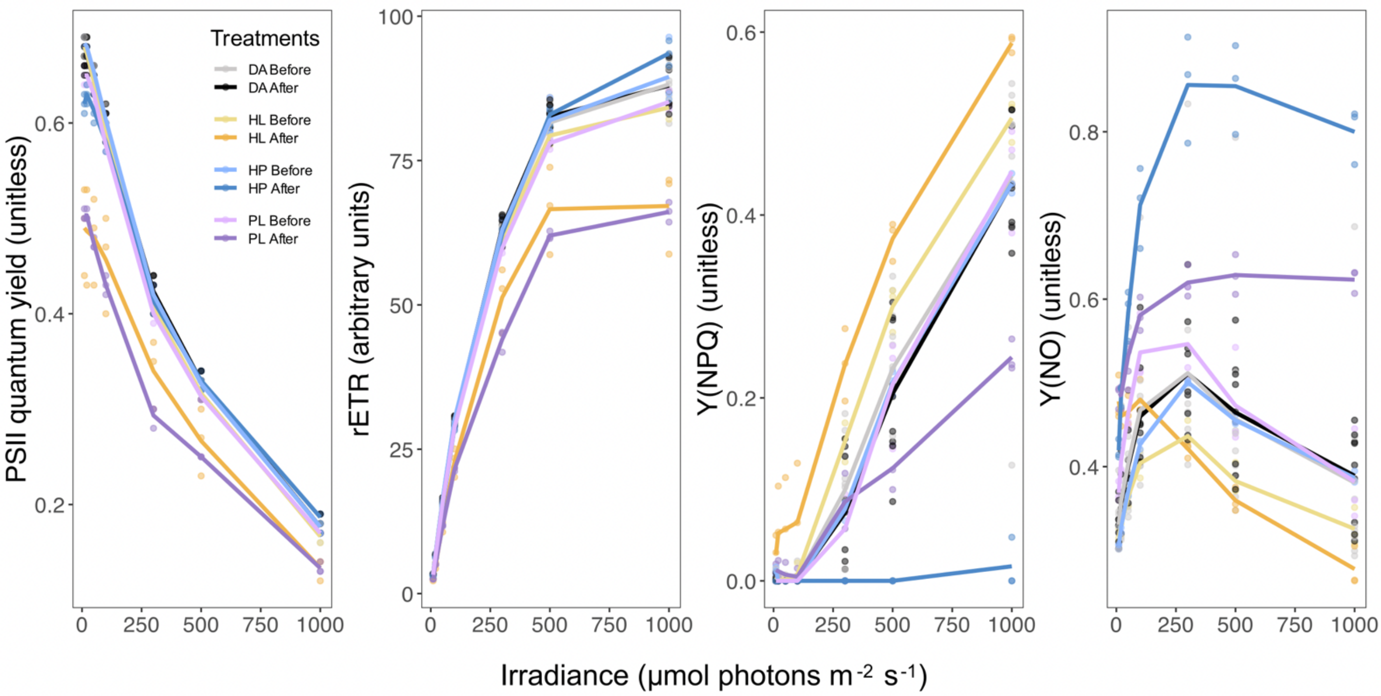


***Figure s3.*** *Photosynthetic efficiency profiles under treatments. Photosystem II (PSII) quantum yield (unitless), rETR (arbitrary unit), Y(NPQ) (unitless), and Y(NO) (unitless) response curves of sediment-associated biofilm community, under the three stresses: High-Light (HL), Hydrogen Peroxide (HP) and Cold Atmospheric Plasma (PL), compared to the Dark-Adapted control samples (DA). All photosynthetic parameters measured with n = 3.*

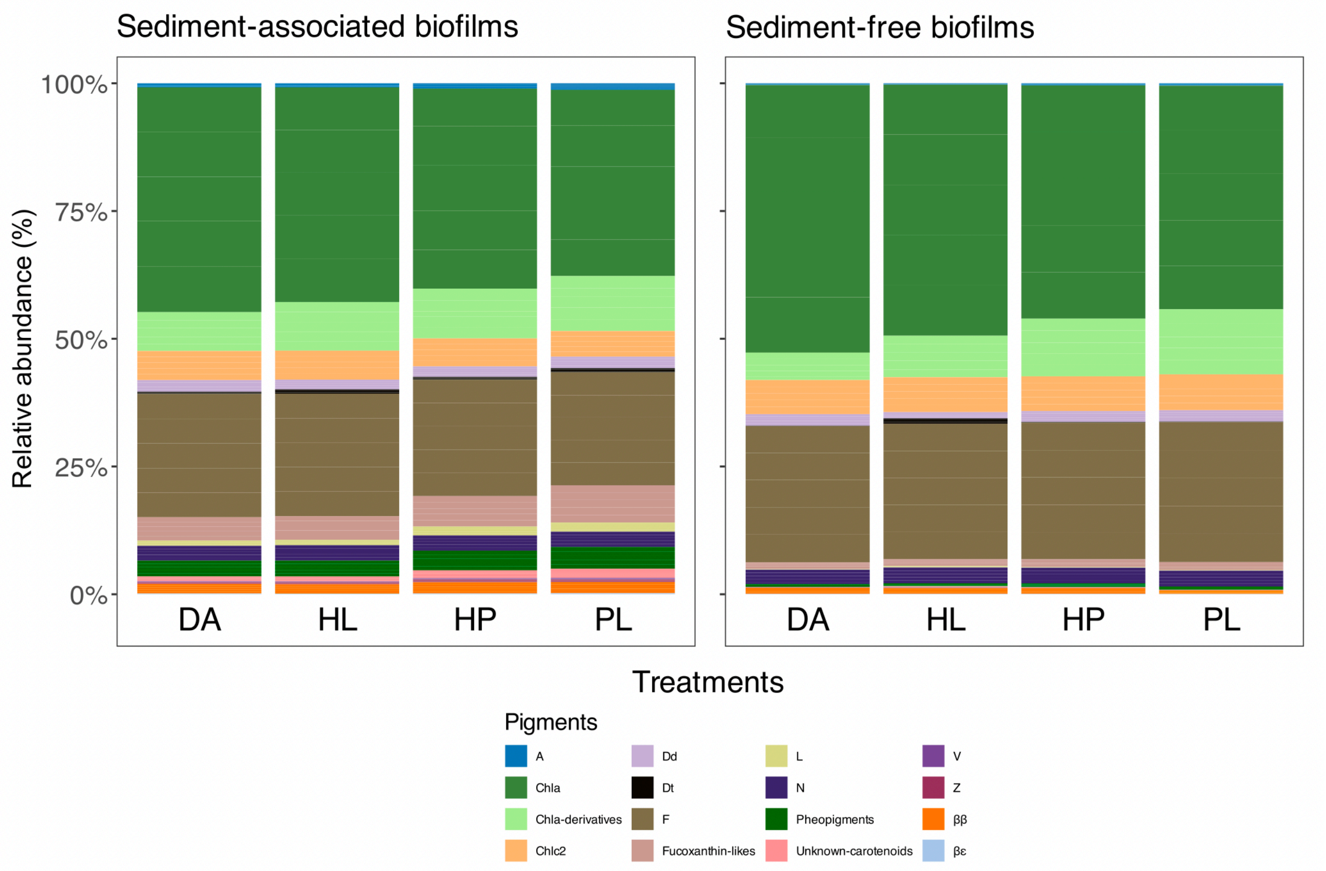


***Figure s4.*** *Barplots of the diversity and relative proportions (%) of the pigment composition of sediment-associated and sediment-free biofilm communities under the applied stresses: High-Light (HL), Hydrogen Peroxide (HP) and Cold Atmospheric Plasma (PL), compared to the Dark-Adapted control samples (DA). Pigments abbreviations are shown in supplementary Table 2.*

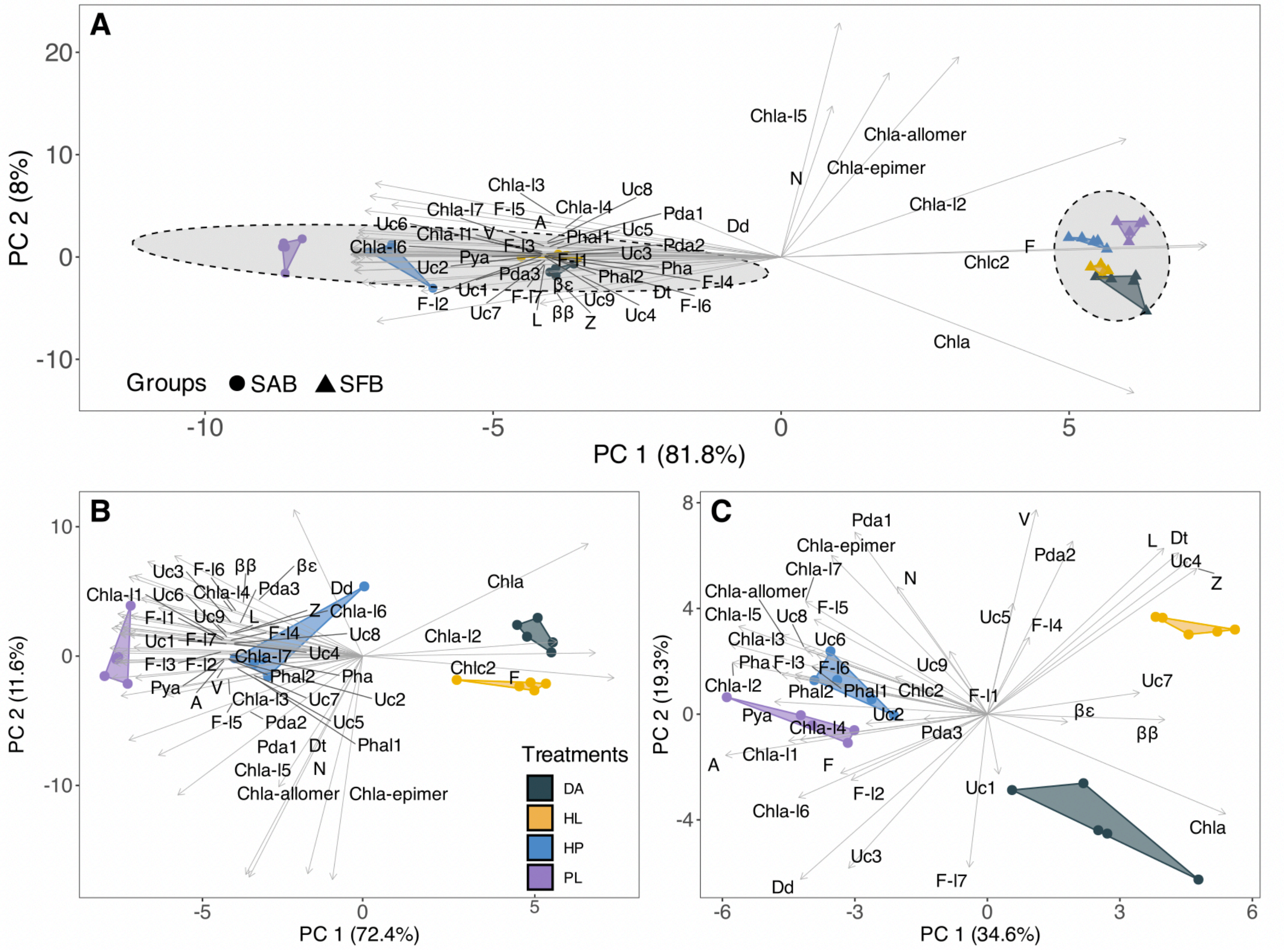

***Figure s5.*** *Biplots of Hellinger-transformed relative pigment proportions and the projection of pigment variables onto the first two principal components (PC1 and PC2) from PCA;* ***A.*** *Samples are grouped by Sediment-Associated Biofilm (SAB) and Sediment-Free Biofilm (SFB) communities, under the applied treatments: Dark-Adapted control (DA), High-Light (HL), Hydrogen Peroxide (HP), and Cold Atmospheric Plasma (PL). Dashed lines indicate 95% confidence ellipses;* ***B.*** *Biplot showing only the sediment-associated biofilm pigment compositions under the applied treatments;* ***C.*** *Biplot showing only the sediment-free biofilm pigment compositions under the applied treatments. The abbreviations for pigment names are defined in supplementary Table 2.*

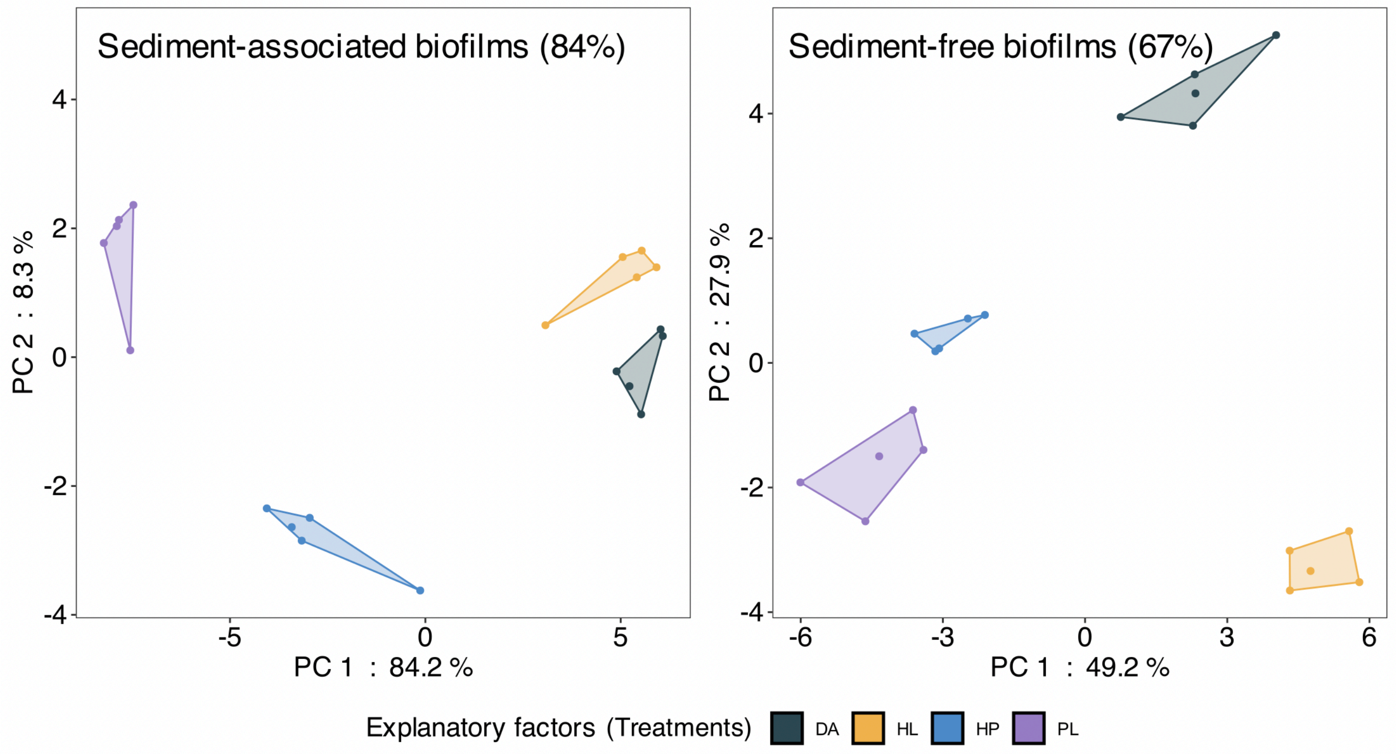


***Figure s6.*** *Between-class analysis based on the first two principal components of the PCA (supplementary Figure 5B and C), performed on pigment composition matrices (%) of sediment-associated and sediment-free biofilms under the applied treatments: Dark-Adapted control (DA), High-Light (HL), Hydrogen Peroxide (HP), and Cold Atmospheric Plasma (PL). Treatments are used as the explanatory grouping variable to account for the inertia observed in the pigment dataset and to reveal the synergy between vertical migration and pigment dynamics in response to the stresses. The role of migration in modulating the pigmentary system is highlighted by the greater separation between DA and HL treatments within the sediment-free biofilm community compared to sediment-associated biofilm samples, indicating that vertical migration in the sediment-associated biofilm samples contributes to mitigate light-induced impact.*

***Table s5.*** *Average xanthophyll concentrations (µg pigments $g^{-1}$ dry matter µ$g^{-1}$ of total Chlorophyll a ± SD) and de-epoxidation state values (DES, %), for both Sediment-Associated Biofilm (SAB) and Sediment-Free Biofilm (SFB) communities, under Dark-Adapted control (DA), High-Light (HL), Hydrogen Peroxide (HP) and Cold Atmospheric Plasma (PL) treatments.*

| Treatment | [Diadinoxanthin] | [Diatoxanthin] | [Diadinoxanthin + Diatoxanthin] | DES% |
|---|---|---|---|---|
| **DA SAB** | 4.2e-02 (4.1e-04) | 8.1e-03 (3.3e-04) | 5.0e-02 (7.1e-04) | 16.4 (0.5) |
| **HL SAB** | 3.5e-02 (9.3e-04) | 1.5e-02 (9.1e-04) | 5.0e-02 (7.4e-04) | 30.4 (1.7) |
| **HP SAB** | 3.9e-02 (5.9e-04) | 1.0e-02 (4.2e-04) | 4.9e-02 (7.9e-04) | 21.3 (0.7) |
| **PL SAB** | 4.4e-02 (1.4e-03) | 1.6e-02 (5.6e-04) | 5.9e-02 (1.8e-03) | 26.5 (0.6) |
| **DA SFB** | 3.9e-02 (1.0e-03) | 1.8e-03 (1.2e-04) | 4.0e-02 (1.1e-03) | 4.4 (0.3) |
| **HL SFB** | 2.1e-02 (1.4e-03) | 1.9e-02 (7.6e-04) | 4.0e-02 (1.2e-03) | 46.5 (2.2) |
| **HP SFB** | 3.7e-02 (1.9e-04) | 2.4e-03 (2.3e-04) | 3.9e-02 (3.2e-04) | 6.1 (0.6) |
| **PL SFB** | 3.9e-02 (6.5e-04) | 1.9e-03 (1.6e-04) | 4.1e-02 (7.1e-04) | 4.7 (0.4) |
