## supplementary materials and methods for "Reactive oxygen species trigger downward vertical migration in diatom microphytobenthic biofilms as a strategy to cope with oxidative stress"

**Microphytobenthos sampling.** A total of 6 biofilm boxes were collected (3 for sediment-associated biofilm samples, 3 for sediment-free biofilm samples). Each box corresponds to the sampling of an area of approximately 0.0625 $m^2$ (25 x 25 cm) from the breeding pond. The seawater used was filtered through a sand filter and subsequently treated with ultraviolet light ($pH_{\text{filtered seawater}}$= 7.9). The analysis of the dry organic matter (dry matter burned at 490°C for 2 hours, *n=3*) for sediment-associated and sediment-free biofilms indicated 19.2 ± 0.4% and 22.7 ± 1.7% organic matter in the dry matter, respectively.

**Oxidative treatments.** The experiment was conducted on May 1st, 2nd, and 3rd, 2024. Due to the time required for triplicate RLC measurements before and after stress exposure for each treatment, as well as the specific procedures involved in the hydrogen peroxide and plasma stress protocols, the experiment had to be distributed across these three days to accommodate the experimental setup. The high-light stress was performed on 01 May 2024, the plasma stress on 02 May 2024, and the hydrogen peroxide on 03 May 2024, with dark-adapted controls samples measured in parallel across all three days.

**Light monitoring.** The MSC15 Gigahertz-Optik spectroradiometer was used to ensure an exposure of 1500 µmol photons $m^{-2}$ $s^{-1}$ under the high-light stress and an ambient irradiance below 5 µmol photons $m^{-2}$ $s^{-1}$ during the dark-adapted control, plasma, and hydrogen peroxide treatments.

**Plasma treatment.** Plasma generation was achieved using TDK CeraPlas piezoelectric direct discharge (PDD) technology, which uses a lead zirconate titanate (LZT) ceramic crystal to ionize atmospheric gas. This process was powered by the PiezoBrush PZ3 device from Relyon Plasma GmbH (8W, 100% power), with the NearField module suited for conductive surfaces. This setup enables the generation of cold atmospheric pressure plasma at low temperatures, producing individual transient thermal plasma filaments. The cold plasma appeared as multiple, unstable micro-discharges in the gaseous phase, leading to the formation of reactive species. Exposure to cold atmospheric plasma produced by the Piezobrush PZ3 device generates a composite stress characterized by slight pH acidification, minimal UV exposure with negligible effects[1], and above all a substantial production of reactive oxygen and nitrogen species[2]. As described in previous work, the plasma generates short-lived radical species (O, OH, NO) as well as longer-lived species ($O_3$, $N_2$*) which, upon contact with the water surface, recombine into more stable compounds such as $H_2O_2$, $NO_2^-$ and $NO_3^-$ that can diffuse into the aqueous phase and reach the diatoms[3]. In accordance with the characterization of the Piezobrush PZ3 source performed previously, RONS constitute the main chemical signature induced by cold atmospheric plasma treatment[2]. To chemically characterize the signature induced by the source on the filtered seawater, immediately after plasma exposure, pH and the concentrations of $H_2O_2$, $NO_3^-$ and $NO_2^-$ were determined using a reflectometer (QUANTOFIX Relax, Macherey-Nagel, Germany). This instrument enables instrumental reading of conventional QUANTOFIX and pH-Fix colorimetric test strips, combining the simplicity of dip-stick assays with the reproducibility of photometric analysis. For pH measurements, two types of pH-Fix test strips were employed: pH-Fix 2.0-9.0 and pH-Fix 6.0-7.7, to obtain a precise resolution around neutrality. Reactive oxygen and nitrogen species were quantified (*n = 3*) using semi-quantitative QUANTOFIX test strips, read by the QUANTOFIX Relax:

- $H_2O_2$: QUANTOFIX Peroxide 25 test strips, instrumental measuring range 0.5-25 mg $L^{-1}$ $H_2O_2$.
- $NO_3^-$: QUANTOFIX Nitrate 250 test strips, instrumental measuring range 4-250 mg $L^{-1}$ $NO_3^-$.
- $NO_2^-$: QUANTOFIX Nitrite 25 test strips (0.5-25 mg $L^{-1}$ $NO_2^-$) and QUANTOFIX Nitrite test strips (0.5-80 mg $L^{-1}$ $NO_2^-$), depending on the expected concentration range.

For each sample, a strip was briefly immersed in the seawater sample, excess liquid was removed, and the strip was inserted into the QUANTOFIX Relax after the specified reaction time. The reflectometer automatically identified the test, read the color fields, and returned concentrations in mg $L^{-1}$. Concentrations of $H_2O_2$, $NO_3^-$ and $NO_2^-$ were subsequently converted to mmol $L^{-1}$ using their respective molar masses.

The characterization shows very little variation in pH (-0.3 units) but a marked increase in $H_2O_2$ (209 µM), $NO_2^-$ (903 µM) and $NO_3^-$ (2.84 mM) after 10 minutes of exposure on filtered seawater (see figure below). This stress is progressive, as RONS concentrations depend on treatment duration (as shown by intermediate values measured at 5 minutes). Altogether, the results indicate a major oxidative impact of plasma treatment driven by the induction of $H_2O_2$. The increase in nitrate does not represent a relevant stressor for diatoms, despite concentrations higher than those typically found in pelagic and sedimentary environments[4, 5]. Indeed, diatoms are capable of accumulating intracellular nitrate concentrations higher than those measured in the treated seawater[4]. Nitrites, on the other hand, remain moderately reactive at neutral pH, with a much slower reaction kinetics with biomolecules compared to hydrogen peroxide. Finally, the filtered seawater exhibited sufficient buffering capacity to maintain a nearly stable pH during exposure. Altogether, the results converge toward a conclusion: the dominant feature of the stress induced by this plasma source is an oxidative stress strongly mediated by the production of $H_2O_2$.

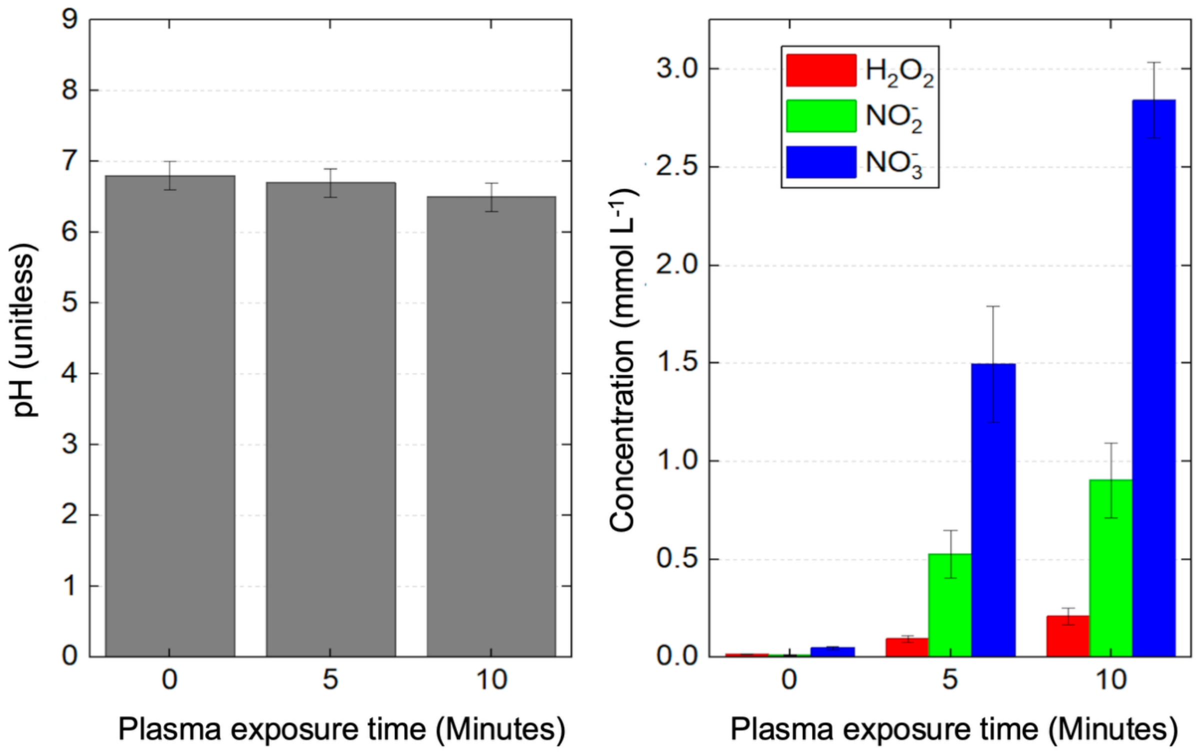


**Hydrogen peroxide treatment.** Based on the $H_2O_2$ plasma generation results, the stress concentration for the hydrogen peroxide treatment was set at 200 µM. This ROS was selected for its long lifespan, high membrane permeability, key role in signaling pathways, ease of use, and cost-effectiveness. The solution was prepared from a stabilized $H_2O_2$ stock solution from Sigma Aldrich, ref. 216763 (water as a carrier; stabilizers: ~0.5 ppm stannate-based compounds and ~1 ppm phosphorus-based compounds; hydrogen peroxide at ~30%). To reach the desired final concentration, 100 µL of the 9.7912 M stock solution were diluted in 10 mL of filtered seawater, yielding a concentrated working solution of 97 912 µM. For the sediment-associated biofilm Petri dishes, 82 µL of this working solution were added to 40 mL of filtered seawater before distributing 4 mL into each replicate Petri dish ($C_{f\ sediment\text{-}associated\ biofilms}$ recalculated = 200.31 µM). For sediment-free biofilms, 8 µL of the working solution were added directly to the 4 mL algal suspension, which was homogenized by repeated pipette aspiration and dispensing ($C_{f\ sediment\text{-}free\ biofilms}$ recalculated = 195.43 µM). For both final concentrations, the stock solution was diluted by a factor of approximately 1:5000.

**18S rRNA gene amplicon sequencing.** The eukaryotic primers TAReuk454FWD1 (5′-CCAGCASCYGCGGTAATTCC-3′) and TAReukREV3 (5′-ACTTTCGTTCTTGATYRA-3′), generate a 369-base amplicon, whereas the prokaryotic primers 338F (5′-

ACTCCTACGGGAGGCAGCA-3′) and 806R (5′-GGACTACHVGGGTWTCTAAT-3′), produce a 468-base amplicon. The use of these universal primer pairs allows broad coverage of both eukaryotic and prokaryotic communities. DNA extraction was performed at the Concarneau Marine Station on the dry matter of the samples using the NucleoSpin Soil Mini Kit. SL1 buffer was used instead of SL2 due to the predominantly mineral nature of the samples, and SW2 was performed with ultrapure 99.8% alcohol. A 50 µL aliquot of the extracted DNA was sent to the Sequencing Platform BMKGENE for amplification of both markers.

The SAMBA workflow relies on the *DADA2*[6], *QIIME2*[7], and *phyloseq*[8] tools. Taxonomic assignment of sequences was performed using a classification algorithm based on a bootstrap approach, with a minimum confidence threshold of 70% required to validate assignments.

Following the taxonomic assignment of eukaryotes using PR2, diatoms of the genus *Gyrosigma* were not identified despite their significant presence observed microscopically in the microphytobenthic biomass. To refine taxonomic identification of diatoms, all unassigned raw ASVs (ATCG sequences) at the genus level were retrieved and manually verified using BLASTn. For *Gyrosigma* assignment, a combination of three decision thresholds was applied (E-value < 0.0001; query coverage = 100%; percentage identity > 97%). Furthermore, a taxonomic assignment was only validated if *Gyrosigma* was the dominant match in the suggested NCBI results list.

**Photosynthetic parameters.** Measurements were performed using the MonitoringPen MP 100-E(v3.2.1.1) and the FluorPen software (v1.0.6.1) was used. The actinic blue light (LED excitation at λ470 nm) and saturating pulses of the MonitoringPen were used for Light curve 3 protocol (10, 20, 50, 100, 300, 500 and 1000 µmol photons $m^{-2}$ $s^{-1}$). Optimal parameters for the superpulse and flashpulse, tested on the microphytobenthic community, were set at 60% (1 800 µmol photons $m^{-2}$ $s^{-1}$) and 30% (900 µmol photons $m^{-2}$ $s^{-1}$), respectively. Photophysiological measurements were conducted in water at a 2 mm distance from microphytobenthic cells. Fluorescence emission was collected between λ667 and 750 nm. A 10-minute period following the high-light exposure is required to ensure stable re-oxidation of the plastoquinone pool in the microphytobenthic community.

**Migration $F_0$.** Biomass tracking based on surface $F_0$ was used to monitor vertical migration responses under stress. Ft measurement required the Flash Pulse, which was set to 100%. The number of replicate measurements and monitoring duration varied, reflecting differences in experimental constraints (5 to 8 replicates per treatment and monitoring periods ranging from 1.5 to 3.5 hours).

**Lipophilic pigments.** The injection method applied to each dry weight sample used the following parameters: $V_{inj}$ = 100 µl, 40-minute program, no TOF and column and sampler temperatures set at 30°C and 14°C (4°C) respectively. Phytoplankton pigment standards from DHI, for calibration curves, were re-evaporated from their original solvents, re-dissolved in 95% MeOH (2% ammonium acetate), and injected using the same method. Calibration curves were determined by linear regression for β-carotene, chlorophyll *a*, chlorophyll *c2*, diadinoxanthin, diatoxanthin, fucoxanthin, lutein, and violaxanthin. Pigments were integrated at their maximum absorption wavelengths, except for chlorophyll *a*, which was integrated at three maximum distinct peaks (λ412, 431, and 663 nm) to resolve overlapping spectra between certain pigments. Chromatogram integration was performed using Agilent Mass Hunter Qualitative Analysis (v10.0).

**Theoretical Qphar.** The pigment concentrations were also used to reconstruct the “typical” absorption spectrum of the microphytobenthic community (dark-adapted sediment-free biofilms undisturbed profiles), following **eq.1** described previously[9]. A correction for the cellular package effect was then applied using equations previously published[10]. This approach allows quantification of the percentage of light attenuation at each wavelength caused by intracellular pigment packing. The corrected absorbance spectrum ($a_{microphytobenthos}^{rec,pac}$; dimensionless), was then weighted by the raw incident irradiance in the calculation of Qphar according to **eq.2** [9]. This procedure provides the best possible estimate of the light actually absorbed by diatoms at the top of substrata during the high-light treatment.

$$a_{mpb}(\lambda) = \sum_{i}^{n} a_i(\lambda) \times C_i \quad \textbf{(1)}$$

$$Qphar(\lambda) = Q(\lambda) - (Q(\lambda) \times e^{-a_{mpb}^{rec,pac}(\lambda)}) \quad \textbf{(2)}$$

*$a_i(\lambda)$ ($m^2\ mg^{-1}$) refers to the concentration-specific absorption spectra of pigments provided in previous work[11]. $C_i$ ($mg\ m^{-2}$) corresponds to the pigment concentrations per petri dish (pigments included in the spectral reconstruction: Antheraxanthin, β-β-carotene, β-Ɛ-carotene, Chlorophyll a, Chlorophyll c2, Diadinoxanthin, Diatoxanthin, Fucoxanthin, Lutein, Neoxanthin, Pheophorbide a, Pheophytin a, Violaxanthin, Zeaxanthin). Q(λ) (µmol photons $m^{-2}\ s^{-1}$) is the raw photon flux density available at the top of the substrata.*

In the context of this study, the estimated Qphar for sediment-free biofilms provides a first-order approximation of the light dose absorbed by diatoms, but should be interpreted with caution, given the numerous simplifications involved. Several limitations arise:

First, pigment modifications occurring during the high-light exposure (*e.g.* changes in the de-epoxidation state) are not taken into account, because the theoretical Qphar under high-light is calculated from the incident high-light irradiance weighted over the dark-adapted sediment-free biofilm samples absorption spectrum. Second, the reconstructed community absorption spectrum is based on pigment-specific absorption measured in solvent[11], whereas in vivo absorption differs substantially. Qphar values are also highly variable because they depend on pigment composition and are therefore species-specific. Additional sources of variability include structural features such as frustule presence and thickness, which modify the incident spectrum before it reaches the pigments[9].

Another major limitation in a microphytobenthic context is the strong pigment packaging effect associated with large diatom cells. Because cell volume scales with packaging, larger diatoms absorb less light per pigment unit. In our biofilm dominated by very large cells (hundreds of µm), neglecting this effect would make Qphar estimates even less realistic. Although we attempted to correct for packaging, this required several additional steps and introduced further approximations, including:

- Approximation of cell biovolume calculated by prism parallelogram-base equation as previously described[12], with average surface measurements (length × width) of the dominant species *P. strigosum* (*n = 30*), and combined with mean cell thickness values[13].
- Equivalent spherical diameter (ESD) biovolume conversion, adding a new approximation.
- Approximating the effective cellular/biofilm height traversed by incident light.
- Extrapolating phytoplankton-based models to large microphytobenthic diatoms, including the use of the absorption coefficient value of microplankton (>20 µm), which is 0.009[10, 14].
- Selection of a single ESD value (average value) within the convergence range of the model proposed previously[10], a realistic and neutral choice, but nonetheless arbitrary.

For sediment-associated biofilms, the calculated Qphar and $Qphar_{package\text{-}corrected}$ values are even less reliable than those for sediment-free biofilm samples. The presence of sediment causes steep attenuation gradients in muddy environments; micrometer-scale vertical migration, changes in cell orientation, or shading by neighboring cells can drastically alter the actual light dose at the cellular level. Sediment and surrounding organic matter also modify the electromagnetic spectrum reaching the cells. It is therefore certain that the irradiance experienced by sediment-free biofilm cells differs substantially from that experienced by sediment-associated biofilm cells, as reflected by the stronger activation of the xanthophyll cycle and the slight increase in Y(NO)m obs. In addition, it is important to note that for sediment-free biofilms, light is received by the cell from all directions, due to the transparency of the Petri dishes.

**Data treatment.** Statistical analyses were performed on RStudio v2025.05.0+496.

### Additional references:


1. Perinban S, Orsat V, Raghavan V. Nonthermal plasma-liquid interactions in food processing: a review. *Compr Rev Food Sci Food Saf* 2019;18:1985–2008. 10.1111/1541-4337.12503
2. Thompson TP et al. Biomedical application of cold plasma: navigating through plasma types and devices by antimicrobial effectiveness and tissue tolerance. *Adv Ther* 2025;8:2400339. 10.1002/adtp.202400339
3. Bruggeman PJ et al. Plasma-liquid interactions: a review and roadmap. *Plasma Sources Sci Technol* 2016;25:053002. 10.1088/0963-0252/25/5/053002
4. Stief P et al. Intracellular nitrate storage by diatoms can be an important nitrogen pool in freshwater and marine ecosystems. *Commun Earth Environ* 2022;3:154. 10.1038/s43247-022-00485-8
5. Rios-Yunes D et al. Sediment resuspension enhances nutrient exchange in intertidal mudflats. *Front Mar Sci* 2023;10:1155386. 10.3389/fmars.2023.1155386
6. Callahan BJ et al. DADA2: high-resolution sample inference from Illumina amplicon data. *Nat Methods* 2016;13:581–583. 10.1038/nmeth.3869
7. Bolyen E et al. Reproducible, interactive, scalable and extensible microbiome data science using QIIME 2. *Nat Biotechnol* 2019;37:852–857. 10.1038/s41587-019-0209-9
8. McMurdie PJ, Holmes S. phyloseq: an R package for reproducible interactive analysis and graphics of microbiome census data. *PLoS ONE* 2013;8:e61217. 10.1371/journal.pone.0061217
9. Prins A et al. Effect of light intensity and light quality on diatom behavioral and physiological photoprotection. *Front Mar Sci* 2020;7:203. 10.3389/fmars.2020.00203

10. Soja-Woźniak M et al. Estimation of the global distribution of phytoplankton light absorption from pigment concentrations. *J Geophys Res Oceans* 2022;127:e2022JC018494. 10.1029/2022JC018494
11. Clementson LA, Wojtasiewicz B. Dataset on the absorption characteristics of extracted phytoplankton pigments. *Data Brief* 2019;24:103875. 10.1016/j.dib.2019.103875
12. Hillebrand H et al. Biovolume calculation for pelagic and benthic microalgae. *J Phycol* 1999;35:403–424. 10.1046/j.1529-8817.1999.3520403.x
13. De Tommasi E et al. Multiple-pathways light modulation in *Pleurosigma strigosum* bi-raphid diatom. *Sci Rep* 2024;14:6476. 10.1038/s41598-024-56206-y
14. Uitz J et al. Vertical distribution of phytoplankton communities in open ocean: an assessment based on surface chlorophyll. *J Geophys Res* 2006;111:C08005. 10.1029/2005JC003207
